# Unlocking the strength of inducible promoters in gram-negative bacteria

**DOI:** 10.1101/2022.04.23.489285

**Authors:** Andrés Felipe Carrillo Rincón, Natalie G. Farny

## Abstract

Inducible promoters, such as the *lac* and *tet* promoters, are ubiquitous biotechnology tools. Inducible bacterial promoters have a consistent architecture including two key elements: the operator region recognized by the transcriptional regulator proteins (e.g., LacI and TetR, and the -10 and -35 consensus sequences required to recruit the sigma (σ) subunits of RNA polymerase to initiate transcription. Despite their widespread use in molecular biology, there remain problems with current inducible expression systems. Leaky transcription in the OFF state remains a particular challenge. Here we have updated the architecture of the *lac* and *tet* expression systems to improve their strength, control, and portability. We modified the genetic architecture of the *lac* and *tet* expression systems to contain consensus -10 and -35 sequence boxes to be strongly targeted by σ^70^, to incorporate of a strong ribosome binding site recognized broadly by gram-negative bacteria, and to independently control of the transcriptional regulators by optimized constitutive promoters. To test the promoters, we use the far-red fluorescent protein mCardinal, which we demonstrate significantly improves the signal-to-background ratio of promoter measurement assays over widely utilized green fluorescent proteins. We validate the improvement in OFF state control and inducibility by demonstrating production of the toxic and aggregate-prone cocaine esterase enzyme CocE. We further demonstrate portability of the promoters to additional gram-negative species *Pseudomonas putida* and *Vibrio natriegens*. Our results represent a significant improvement over existing protein expression systems that will enable advances in protein production for various biotechnology applications.

**Significance:** Many of the latest advances in pharmaceuticals, materials, and foods involve the production of recombinant proteins from bacterial hosts. However, the regulated production of enzymes and functional protein products that are toxic to their microbial hosts remains a challenge. Our work provides new tools that enable tight control over expression of protein products in bacterial host strains. We show that our tools function not only in the broadly utilized *Escherichia coli*, but also in other gram-negative bacteria including the soil organism *Pseudomonas putida* and the marine bacterium *Vibrio natriegens*. Our technology will facilitate more efficient production of a broader range of protein products in diverse microbial hosts.

## Introduction

Inducible promoters, such as the *lac* and *tet* promoters and their derivatives [1, 2], are workhorses of the scientific community. Inducible bacterial promoters have a consistent architecture including two key elements: the operator region recognized by the transcriptional regulator proteins (e.g., LacI and TetR [3-5], and the -10 and -35 consensus sequences required to recruit the sigma (σ) subunits of RNA polymerase to initiate transcription [6]. Over the years, there have been several notable improvements to the lac promoter to increase its strength and improve its regulation. The -10 and -35 boxes of the original *lac* promoter (Fig. 1Ai) were updated to generate improved variants such as the *lacUV5* (Fig. 1Aii) and *tacI* (Fig. 1Aiii) promoters [7, 8]. In the *lacUV5* promoter, the -10 box of the original *lac* promoter was replaced by the Pribnow box (TATAAT) [9]. Later the *lacUV5* and *trp* promoters were combined to create the *tacI* promoter, which increased transcription 11-fold compared to its predecessor with the incorporation of the -35 consensus box TTGACA [7] [10]. The *tet* promoter (Fig. 1Bi) shares the highly conserved -35 hexamer with *tac* promoter, however it does not contain the Pribnow box.

**Figure 1.**
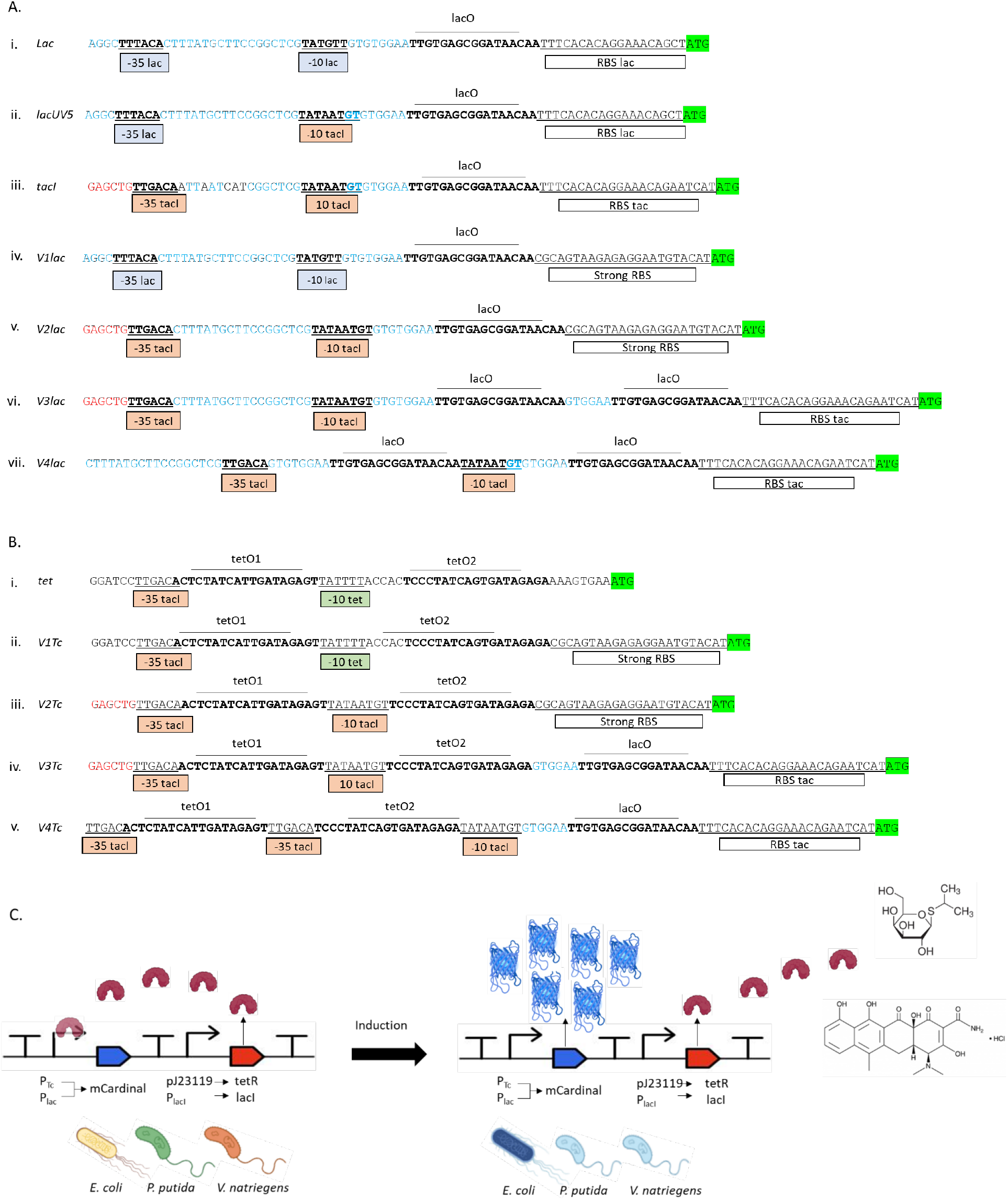
Architecture of the synthetic of *lac* and *tet* promoters. The sequences of the operators lacO and tetO are overlined, in bold text. The -35 and -10 sequences of the promoters are underlined. Blue and green boxes represent the native -35 and -10 sequences, while orange boxes show the consensus σ^70^ -35 and -10 box sequences. Blue nucleotides indicate conserved regions of the original *lac* promoter. **A**. Original *lac* (i) and synthetic *lac* (ii-vii) promoters. **B**. Original *tet* (i) and synthetic *tet* (ii-v) promoters. **C**. Schematic representation of the function of synthetic *lac* and *tet* promoters. Each transcriptional unit is insulated by terminators. LacI/TetR regulators bind to the operators lacO/tetO. Addition of the inducers IPTG or anhydrotetracycline (aTc) remove the repressor allowing transcription from the promoters.

Despite their ubiquitous use in molecular biology, there remain problems with the current *lac*-based inducible expression systems. Leaky transcription in the OFF state remains a consistent challenge [11]. Tight transcriptional control is indispensable to produce high yields of challenging recombinant proteins, including toxic genes and proteins that are difficult to process or fold for a heterologous host [12]. Various strategies have been applied to avoid leakage and amplify the dynamic range of the inducible expression systems [13]. For example, in the widely utilized pET system, target proteins are driven indirectly by controlling expression of the T7 polymerase under the *lacUV5* promoter, and then driving the target gene transcription by the T7 promoter. Still, low basal level T7 transcription leads to leaky expression of the target gene. To combat this problem further, *E. coli* strains such as pLysS and pLysE with integration of the T7 lysozyme that inhibits low level T7 activity are used to obtain tighter transcriptional control. However, a host strain with T7 RNA polymerase under control of the *lacUV5* promoter, and integration of T7 lysozyme, are required to tightly regulate target gene expression [14].

Expression with the *tet* system has historically provided tighter transcriptional control than *lac* derived promoters, does not required specialized strains (such as the BL21), and full induction is achieved by anhydrotetracycline (aTc) at concentrations that do not cause growth defects due the high affinity of aTc to TetR, and its imperceptible antibiotic activity [15, 16]. Remarkably, in the uninduced state just one mRNA molecule per three cells is produced, and up to 5000-fold induction has been reported [15]. However, the yields of recombinant protein obtained by the tet expression system remains low compared to the pET expression system when using *E. coli* as heterologous host.

The repertoire of organisms used both in academic and industrial settings is rapidly expanding. To address challenges related to complex protein expression in *E. coli*, other chassis organisms such as *Pseudomonas putida* and *Vibrio natriegens* have been employed to produce challenging proteins in a variety of biotechnological processes [17-19]. Both *lac* and *tet* expressions systems have been adapted to *P. putida* and *V. natriegens* [20-22], with varying levels of success, and typically lower total protein yield than achieved in pET. There is currently no universal expression system than can be directly ported between different gram-negative species, yield high quantities of recombinant protein comparable to pET, *and* maintain tight transcriptional repression in the uninduced state.

In this study we have updated the architecture of the *lac* and *tet* expression systems to improve their strength, control, and portability. We modified the genetic architecture of the *lac* and *tet* expression systems in three ways: (1) addition of the consensus -10 and -35 sequence boxes to be strongly targeted by σ^70^, (2) incorporation of a strong ribosome binding site recognized by a broad spectrum of gram-negative bacteria, and (3) independent control of the transcriptional regulators by appropriately-tuned constitutive promoters. We validated the results with the reporter protein mCardinal [23], which significantly improves the dynamic range of promoter measurements over more commonly used green fluorescent proteins. Additionally, we confirmed the advantage of our expression system against the pET system with the production of the cocaine esterase CocE [24], a thermosensitive enzyme capable of metabolizing cocaine into benzoic acid. CocE, which is prone to form inclusion bodies in leaky *E. coli* expression systems, is expressed as a soluble protein using our promoters. Our observations confirm that the expression system presented in this study is a significant improvement over available expression systems in providing tight OFF state control while achieving high yields of recombinant protein, and with the advantage of direct portability to alternative host species.

## Results

### Design of the synthetic lac and tet promoters

The original *lac* promoter (Fig. 1Ai) lacks the -35 conserved box recognized by the σ^70^ subunit of RNA polymerase [25]. Original *lac* and *tet* (Fig. 1Bi) promoters lack the Pribnow (−10) box [15, 25]. We constructed three synthetic variants of each promoter containing the highly conserved σ^70^ consensus hexamers at positions -10 and -35. Version 1 (*V1lac*, Fig. 1Aiv, and *V1Tc*, Fig. 1Bii) of each promoter resembles the original *lac* and *tet* promoters, with the incorporation of a strong RBS from *P. putida* [26]. The version 2 (*V2lac*, Fig. 1Av, and *V2Tc*, Fig. 1Biii) replaces the -35 and -10 boxes of Version 1 for the highly conserved σ^70^ consensus hexamers TTGACA and TATAAT, respectively. Version 3 (*V3lac*, Fig. 1Avi, and *V3Tc*, Fig. 1Biv) incorporates an additional lacO operator and the RBS of the *tacI* promoter. Version 4 (*V4lac*, Fig. 1Avii, and *V4Tc*, Fig. 1Bv) displaces the location of the -10 and -35 boxes (*V4lac*) and incorporates an additional -35 box in *V4Tc*. The synthetic promoters were cloned into the MCS of the shuttle vector pJH0204 (Supplementary Fig. S1) containing the origin of replication pColE1 with 25-30 copies per cell in *E. coli* [27]. The design allowed the incorporation of the transcriptional regulatory genes lacI and tetR after a terminator and under the control of the constitutive promoter pJ23119 (Fig. 1C).

Despite some reports indicating that both LacI and TetR are toxic at high concentrations [28, 29], we did not observe an aberrant phenotype in the *E. coli* strains carrying the pJ23119-tetR construct. However, *E. coli* failed to maintain the pJ23319-lacI construct. The plasmids containing the lacI repressor in this configuration produced slow-growing colonies and yielded plasmids with aberrant restriction patterns, indicating gross plasmid rearrangements. Therefore, we replaced the strong pJ23119 promoter for the native lacI promoter and observed no further toxicity.

The reporter protein mCardinal was incorporated downstream of the synthetic promoters and the plasmids were transformed into *E. coli* DH10B, *P. putida* AG4775 and *V. natriegens* Vmax *X2* strains. The *V2lac-mCardinal* constructs could not be maintained in *E. coli*, likely due the high strength of the *V2lac* promoter. To avoid this abnormal phenotype, the genetic circuit *V2lac-mCardinal* was produced in the pJH0228 vector containing the CloDF13 origin of replication at about 10 copies per cell [30], allowing stable maintenance of the *V2lac* promoter.

### Selection of the fluorescent reporter system

GFP and other green fluorescent protein derivatives are widely used as reporters in bacterial expression systems. However, the intrinsic green fluorescence produced by endogenous molecules, such as proteins containing aromatic amino acids, negatively impact the interpretation of the exogenous green fluorescence generated by the reporter system [31]. The autofluorescence is exacerbated in *P. putida*, a member of the fluorescent *Pseudomonas* species. Under iron limited conditions *P. putida* secretes the siderophore pyoverdine, a soluble fluorescent yellow green pigment [32-34]. Therefore, we explored the advantage of using Far-red fluorescent proteins, specifically mCardinal, a monomeric far red shifted derivative of mKate [35], as wavelengths between 600 and 900 nm are not absorbed by cells and organic molecules [36], thus reducing the noise of endogenous autofluorescence.

To quantify the impact of autofluorescence on the selected gram-negatives strains, we measured the endogenous fluorescence of these strains cultivated in LB over time. All three species emit fluorescence in the green spectrum, as expected (Fig 2A, bars). This fluorescence is constant over time with *E. coli*, however *P. putida* and *V. natriegens* show a tendency to increase the production of molecules that absorb green light at high cell densities (Fig. 2A, bars). In contrast, when measured in the far-red spectrum specific for mCardinal, there was approximately 400 times less detection of autofluorescence (in arbitrary units on the same instrument) in all strains and LB controls as compared to the measurements in the green spectrum (Fig. 2A, lines).

**Figure 2.**
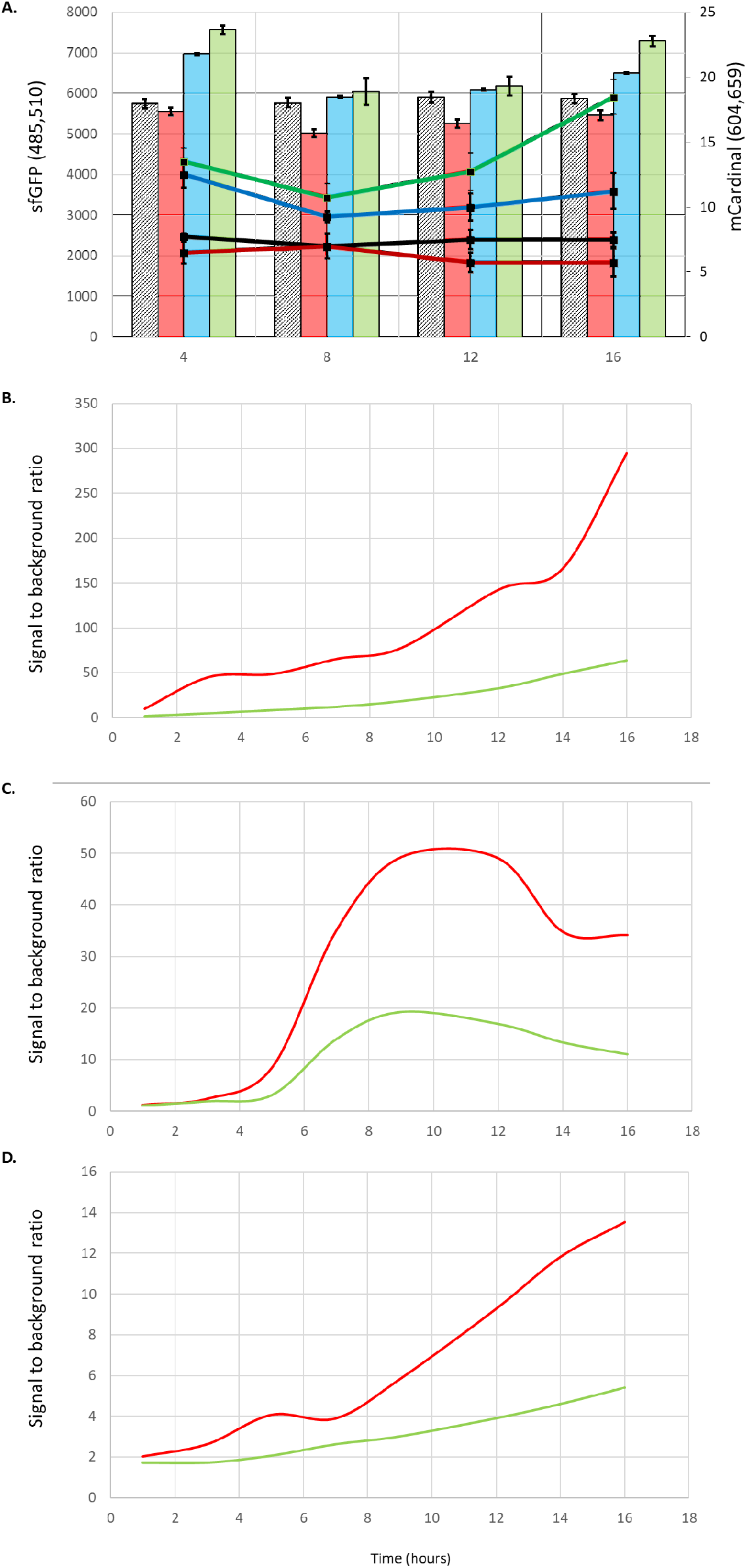
Comparison of sfGFP and mCardinal emissions in gram-negative bacteria. **A.** Bars indicate absolute values of fluorescence measured with sfGFP (excitation 485, emission 510), and lines indicate fluorescence measured with mCardinal (excitation 604, emission 659) of LB alone (Black), *E. coli* DH10B (red), *V. natriegens* (blue) and *P. putida* (green). **B, C & D**. Fluorescence signal-to-background ratio of recombinant strains expressing sfGFP (green) and mCardinal (red) with constitutive *tacI* promoter vs. wild type strains of (**B**) *E. coli* (**C**) *P. putida &* (**D**) *V. natriegens*. N=3. Error bars +/-SD.

To further validate the benefit of using mCardinal instead of sfGFP, we quantified the inherent noise of each reporter system by measuring the fluorescence signal-to-background ratio of both reporter systems expressed under the control of the constitutive *tacI* promoter in our three strains. After 16 hours of growth, the recombinant *E. coli, P. putida* and *V. natriegens* expressing sfGFP emitted 64, 11 and 5 times more green fluorescence than the wild type (not expressing a fluorescent protein), respectively. Meanwhile, mCardinal strains displayed 294, 34 and 13-fold higher red-light emission in *E. coli, P. putida* and *V. natriegens* compared to their wild type controls (Fig. 2B-D). Overall, the use of mCardinal rather than sfGFP significantly improves the dynamic range and facilitates measurement of promoter strength, as there is little endogenous autofluorescence in the far-red spectrum.

### Strength and regulation of synthetic *lac* promoters in *Escherichia coli*

The *lac* promoter, and its derivates, are constitutively active in the absence of its transcriptional regulator lacI. Therefore we first evaluated the strength of the synthetic *lac* promoters without lacI and compared these against the strong *tacI* promoter, which drives high levels of transcription and can result in recombinant protein production of up to 30% of total protein [14]. The *V1Lac* promoter was 10-fold weaker than the *tacI* promoter (Fig. 3A), consistent with previous data observing an 11-fold difference between these two promoters [10]. The *V3lac* promoter matched the *tacI* promoter strength, while the *V4lac* promoter showed 5 times less mCardinal than *tacI* (Fig. 3A). The *V2lac* promoter had to be tested in the low copy plasmid pCloDF13 because the medium copy pColE1 derived plasmid could not be maintained in *E. coli*. Even in the low copy configuration, the *V2Lac* promoter surpassed by ∼1.4-fold the *tacI* promoter and was therefore the strongest constitutive promoter despite its expression in a low copy plasmid. This result suggests that its maintenance in a medium copy plasmid exceeds the sustainable metabolic burden of *E. coli* (Fig. 3A). These results indicate that the incorporation of σ^70^ consensus sequences at positions -35 and -10 efficiently increase the strength of the *lac* promoter.

**Figure 3.**
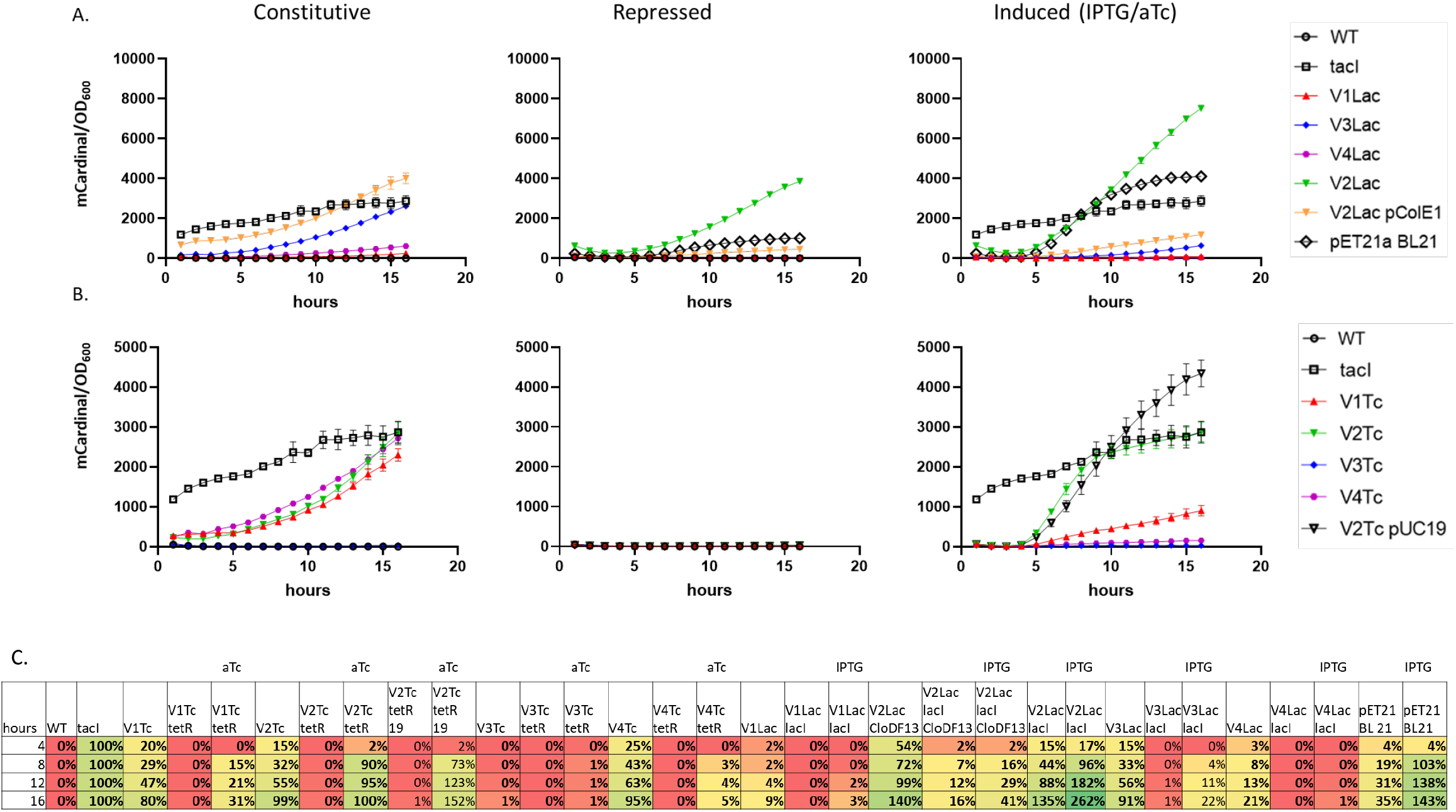
Time course of mCardinal production in *E. coli* DH10B with synthetic *lac* and *tet* promoters. **A.** The synthetic *lac* promoters, and **B**. the synthetic *tet* promoters, with and without the transcriptional regulators tetR and lacI and induced after 3 hours with aTc or IPTG, respectively. **C**. Direct comparison of each promoter under evaluation against the strong constitutive promoter *tacI*. The recombinant DH10B carrying *tacI-mCardinal* was normalized to 100% mCardinal production and the wild type DH10B strain normalized to 0% mCardinal production. For all samples, the fluorescence mean was normalized by the cell density (OD_600_). N=4. Error bars +/-SD.

Next, the transcriptional regulator lacI was incorporated into these circuits to quantify the efficiency of the OFF state. The *V1Lac* and *V4lac* constructs containing the lacI repressor produced a red fluorescence signal comparable to wild type *E. coli* (Fig. 3A), indicating tight transcriptional repression. The *V3lac* promoter could not be totally repressed by lacI, showing a slight tendency to leak, even with the presence of two lacO operators. The *V2lac* promoter with lacI was stably maintained by *E. coli* in both pColE1 and CloDF13-derived vectors. However, the repressed state of the *V2lac* promoter in the low and medium copy plasmids showed rather a constitutive behavior. LacI repressed ∼9-fold the *V2Lac* promoter in the CloDF13 derived plasmid as compared to its constitutive construct (Fig. 3A), while the medium copy version of the *V2lac-lacI* system (which was not sustainable as a constitutive circuit) exhibited a strong constitutive expression ∼1.3-fold higher than the *tacI* promoter, thus showing that one lacO operator region is not sufficient to block transcription of *V2Lac* containing both σ^70^ consensus sequences by LacI, but incorporation of a second lacO reduces the leakage as observed in the *V3Lac* promoter (Fig. 3A).

To verify the inducibility of the synthetic *lac* promoters, *E. coli* was exposed to IPTG at the beginning of exponential phase, at t=3 hours. The *V1lac* and *V4lac* promoters showed minimal induction of mCardinal (Fig. 3A). The *V3lac* promoter, which mimics the *V2lac* promoter with an additional lacO operator, demonstrated a dynamic range of ∼17-fold versus the uninduced state, but could only produce 24% of its full constitutive potential (Fig. 3A). The *V2lac* promoter in the low copy plasmid produced ∼2-fold more mCardinal upon induction; similar behavior was observed in the medium copy version, where production of mCardinal reached the maximum values of all promoters tested at ∼2.6-fold stronger than the *tacI* promoter (Fig. 3A). Overall, our results reveal that incorporation of σ^70^ consensus sequences to the lac promoter improve its strength (*V2Lac*), producing more mCardinal than the strong *tacI* promoter in both, the constitutive LacI-less version in the low copy plasmid and the repressed and induced states in the medium copy plasmid. However, LacI could not repress the transcription of the synthetic *V2Lac* promoter. Thus, additional lacO operators are necessary to turn the lac promoter OFF containing the perfect σ^70^ boxes, as observed in *V3Lac*, though full induction then becomes impossible in this architecture.

### Strength and regulation of synthetic *tet* promoters in *Escherichia coli*

The *tet* promoter also drives transcription constitutively without TetR, thus we evaluated the expression profile of the constitutive synthetic *tet* promoters and compared them against the *tacI* promoter. As observed for the *lac* promoter, replacement of the −10 and −35 sequences with the σ^70^ consensus boxes in the *tet* promoter also increased the constitutive efficiency of the *V2Tc* promoter over the original *V1Tc*, in this case by 20%, and reached the yields obtained with *tacI* promoter (Fig. 3B). The *V4Tc* promoter showed similar results to the *V2Tc*, while the *V3Tc* lost any capability to initiate transcription (Fig. 3B). Further, incorporation of the tetR repressor gene to each of the four circuits under the control of the strong constitutive PJ23119 promoter completely silenced mCardinal production from all the synthetic *tet* promoters, even in the strong *V2Tc* promoter which harbors the optimal combination of consensus sequences recognized by the σ^70^ (Fig. 3B).

To confirm the induction of each promoter and determine its functional dynamic range, we induced each circuit with anhydrotetracycline (aTc). The original tet promoter (*V1Tc*) only achieved ∼37% of its full constitutive potential, reaching only 30% of mCardinal compared to *tacI* promoter confirming the middle level strength of the *tet* promoter (Fig. 3B). The *V2Tc* promoter achieved maximal induction reaching the full potential dynamic range and produced similar yields as the strong *tacI* promoter (Fig. 3B). Interestingly, the *V4Tc* promoter could not be de-repressed by aTc and showed a weak induction of only 5%, and as expected, the *V3Tc* promoter showed no activity (Fig. 3B). Together these results confirm that in *E. coli* the incorporation of the consensus boxes targeted by the σ^70^ is key to boost the expression profile of promoters recognized by this transcription initiation factor and achieve the maximal transcriptional capacity of these promoters upon induction.

### Direct comparison of synthetic *lac* and *tet* expression systems to pET in *Escherichia coli*

In *E. coli*, the pET series of expression plasmids are the most popular systems for recombinant protein production [1]. Therefore, we compared the efficiency of the synthetic *lac* and *tet* promoters directly against pET21a. The pET expression system in the BL21 strain can produce more than 50% of the target gene as the total protein per cell [37], thus we measured mCardinal production in BL21. As expected, the pET system induced by IPTG produced ∼1.5-fold more mCardinal than the strong constitutive *tacI* promoter, which is expected to accumulate up to 30% of total cell protein [14]. Despite the massive production of mCardinal by the pET system, its OFF or uninduced state can be considered as a medium high constitutive expression system, yielding 30% of the constitutive *tacI* promoter as evidenced by mCardinal production (Fig. 3B,C). This transcriptional leakage in the pET system is well known in the scientific community [14].

Among the synthetic *lac* and *tet* promoters, all tested in *E. coli* DH10B, only the *V2Lac* promoter surpassed the pET system in the medium copy plasmid pColE1. The IPTG induced *V2Lac* expression system produced ∼1.8 times more recombinant protein than the pET system, however, the leakage of *V2Lac* promoter equals the induced state of the pET system (Fig. 3A, Supplementary Fig. S2, S3), therefore cancelling the advantages of an inducible expression system. The *V3Lac* expression system showed a tighter control of the OFF state, therefore to increase its strength we inserted the *V3Lac-lacI* construct containing mCardinal into the pUC19 vector, thus increasing the copy number from 30 to 250 copies per cell [38]. *E. coli* failed to maintain the *V3lac-lacI* construct in the pUC19 vector, likely due to the toxicity of LacI. Consequently, we decided to evaluate the *V2Tc-tetR* expression system in the pUC19 vector. *E. coli* stably maintained the *V2Tc-tetR* construct and no phenotypic or genotypic abnormalities were observed as in pUC19-*V3Lac-lacI*. mCardinal production by pUC19-*V2Tc-teR* matched the pET21a expression system, with one exceptional difference, the pUC19-*V2Tc-tetR* maintained complete repression in the uninduced after 12 hours (Fig. 3 B and Supplementary Fig. S2). Thus, pUC19*-V2Tc-tetR* shows significant improvement in transcriptional control and dynamic range over pET21a. Further, equivalent protein production and tighter transcriptional control were achieved without the obligatory use of the BL21(DE3) strain, suggesting our system will be readily portable to other strains and species.

### Expression of the cocaine esterase CocE and production of benzoic acid

To further validate the advantage of our expression system over the pET expression system we evaluated the production of a functional protein product, the cocaine esterase CocE [24]. This enzyme hydrolyses cocaine into benzoic acid and could expand the use of the narcotic compound as raw source in the production of the carboxylic acid widely used as precursor and preservative in the food and pharmaceutical industries. Expression of CocE has previously been shown to be a difficult and laborious task with the pET expression system, because CocE forms inclusion bodies; consequently, long incubation methods at low temperatures are required to isolate sufficient yields of functional CocE [39]. *E. coli* could not support CocE expression in the repressor-less variants of the synthetic promoters, thus confirming this is a toxic protein for *E. coli* (our unpublished observations). Therefore, we evaluated CocE in the *V2Tc-tetR* expression system in both the medium and high copy plasmids, and compared against the pET expression system for the production of benzoic acid. Benzoic acid production is an indication that CocE was correctly folded and properly hydrolyzing cocaine. The soluble fraction recovered from the recombinant *E. coli* strain containing the pET-cocE expression system showed benzoic acid production in the presence of cocaine in the uninduced and induced states, with a clear tendency to form inclusion bodies, as evidenced by the accumulation of protein in the insoluble fraction, upon activation by IPTG (Fig. 4A). The soluble fractions from the recombinant *E. coli V2Tc-tetR-*cocE expression system in both, the medium and high copy plasmid, showed no benzoic acid production in the absence of the inducer, and addition of aTc triggered production of CocE. In both plasmids, benzoic acid production was observed after only one hour of induction (Fig. 4B). However, the high copy plasmid pUC19 underperformed the medium copy plasmid probably due its high strength, leading to the formation of insoluble and inactive forms of CocE. The medium copy plasmid reached the highest benzoic acid production after 3 hours of induction, half the time required in the pET expression system, thus indicating better maturation of the enzyme to perform the breakdown of the cocaine into benzoic acid (Fig. 4B). Together these results confirmed the advantage of the incorporation of the σ^70^ boxes to the tetracycline promoter, keeping the toxic enzyme CocE OFF in the uninduced state and producing sufficient amounts of mature CocE after 3 hours of induction, thus shortening the purification process from 16 hours previously reported [39] to ∼4 hours of fully functional CocE. Our method thus facilitates the scalability of this bioprocess which has potential as an alternative, environmentally friendly method to obtain benzoic by replacing petroleum-based starting materials [40].

**Figure 4.**
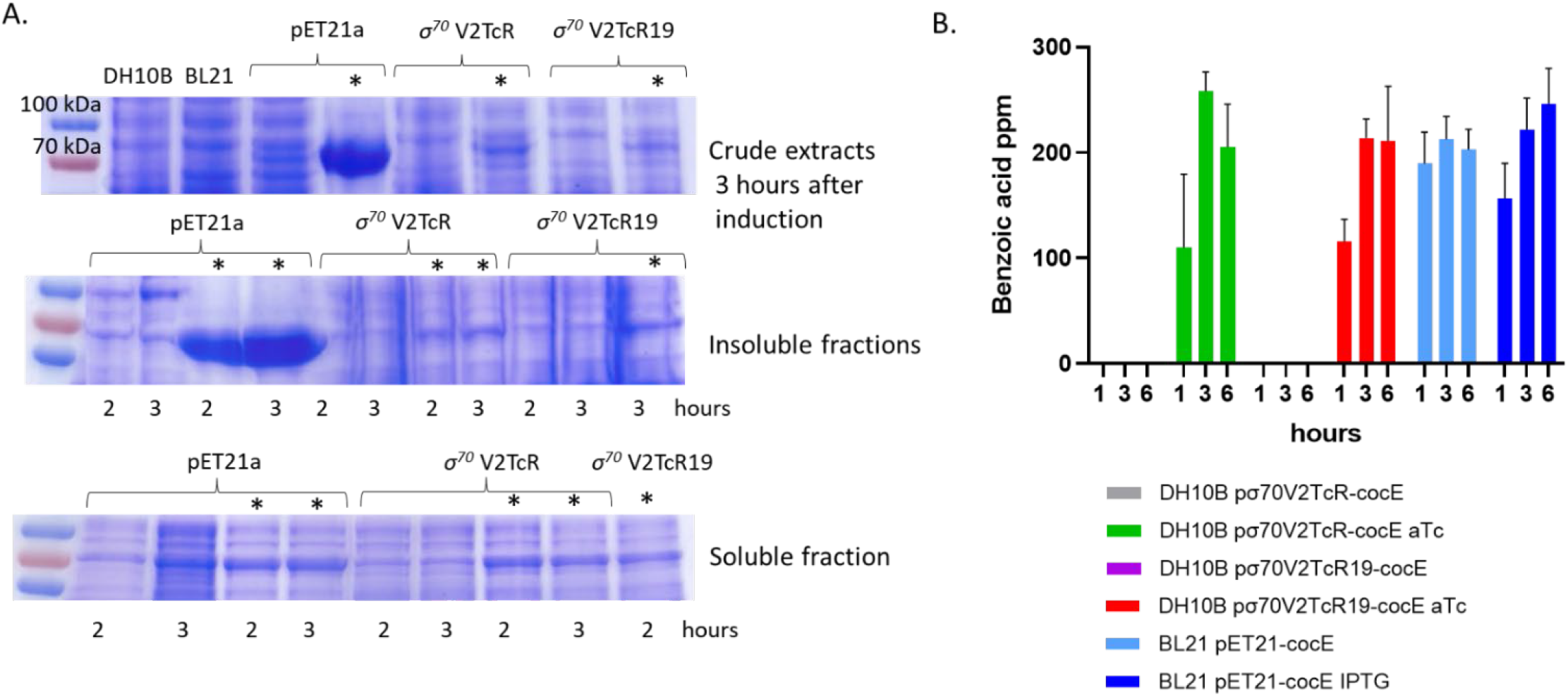
Expression profile of CocE in *E. coli*. **A.** *E. coli* expressing the pET and *V2Tc-tetR* expression systems, uninduced and induced (indicated with an asterisk). **B**. Benzoic acid production by CocE present in the soluble fraction, using cocaine as substrate. N=3. Error bars +/-SD.

### Strength and regulation of synthetic *lac* and *tet* promoters in *Pseudomonas putida*

Genetic control in the soil-dwelling species *P. putida* is of great interest for biotechnological applications due to the ability of this microorganism to synthesize complex natural products and metabolize a variety of xenobiotic compounds [19]. Therefore, we characterized the constitutive strength, repression, and inducibility of our synthetic promoters in *P. putida*. Prior reports in *P. putida* indicated the *tacI* promoter is 40 times more efficient than the *lac* promoter, using mNeonGreen as a reporter [41]. However, the lac promoter systems are reported to have poor dynamic range and instability in high-copy plasmids when harbored in *P. putida* as replicative plasmids [22]. To address this problem, we integrated our genetic circuits into the *P. putida* chromosome [41]. We measured the constitutive strength of *V1lac, V2lac, V3lac* and *V4lac* promoters using mCardinal, and found out that *tacI* is stronger than all the synthetic *lac* variants by ∼3, 1.5, 10 and 33-fold respectively (Fig. 5C). The incorporation of the transcriptional regulator lacI efficiently repressed the *V1lac, V3lac* and *V4lac* promoters, but as seen in *E. coli*, the *V2lac* showed a tendency to leak, though not as egregiously as in *E. coli* (Fig. 5A). Induction by IPTG was barely detected in the *V1lac, V3lac* and *V4Lac* promoters, thus indicating the weakness of the original *lac* promoter in *P. putida* (Fig. 5A). Interestingly and analogously to *E. coli*, the *V2lac* promoter showed ∼27-fold increase in mCardinal production against the uninduced state and surpassed its theoretical dynamic range as compared to the constitutive version, ultimately achieving similar yields as the strong *tacI* promoter (Fig. 5C). These results confirm that the incorporation of the σ^70^ consensus sequences to the *lac* promoter (*V2Lac*) significantly improve the strength of this promoter in the *P. putida* host.

**Figure 5.**
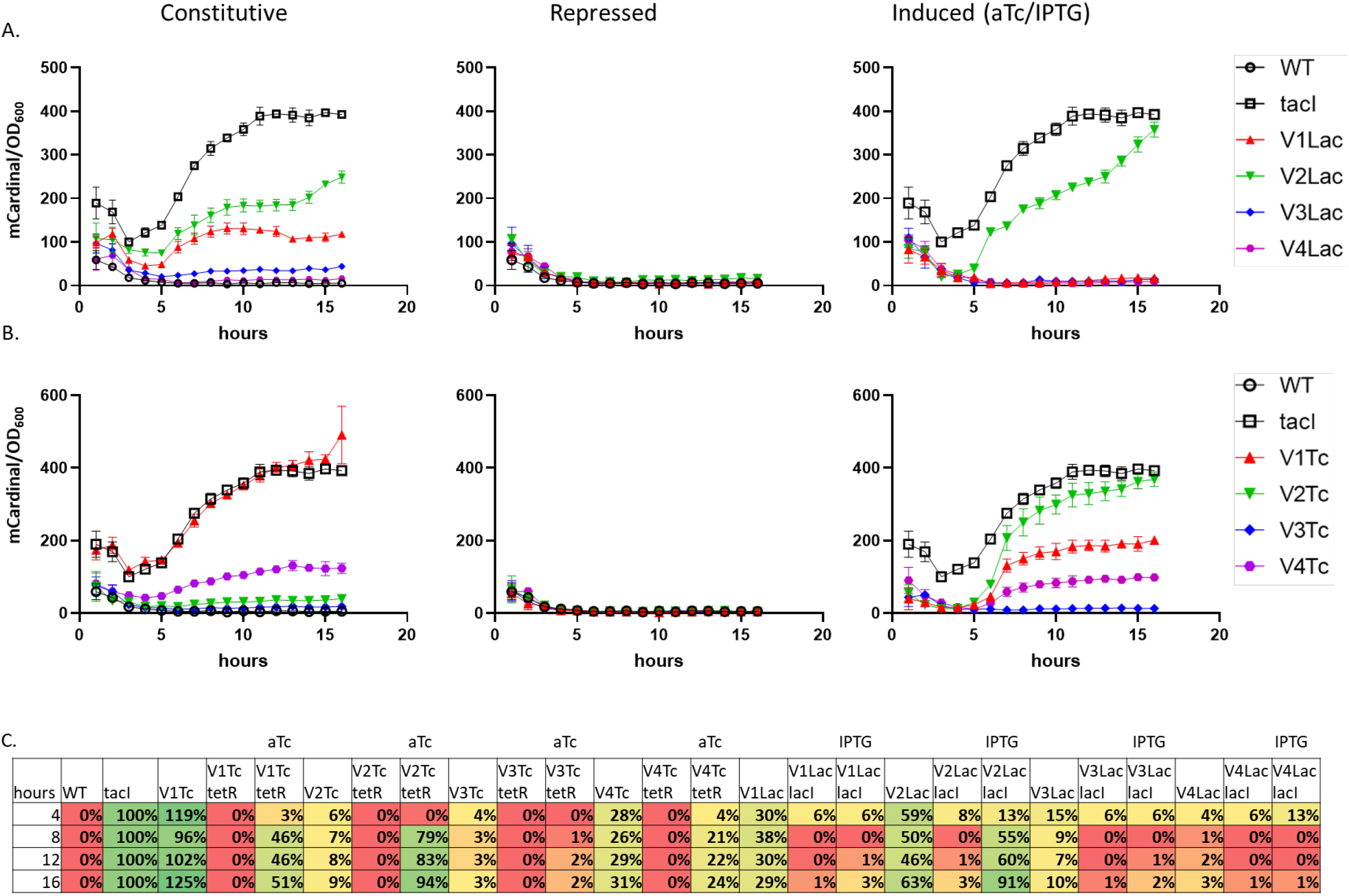
Time course of mCardinal production in *P. putida*. Synthetic *lac* (**A**) and *tet* (**B**) promoters in their constitutive, repressed and induced states. All constructs were integrated in a single copy into the same genomic locus. **(C)** Direct comparison of each promoter under evaluation against the constitutive promoter *tacI* in *P. putida. P. putida* with a single-copy integration of *tacI-mCardinal* was normalized to 100% mCardinal production and the wild type *P. putida* strain normalized to 0% mCardinal production. For all samples, the fluorescence mean was normalized by the cell density (OD_600_). N=4. Error bars +/-SD.

We similarly evaluated the *tet* promoters in *P. putida*. We demonstrated that the original *tet* promoter with a strong RBS (*V1Tc*) is the strongest among all synthetic *tet* promoters in the absence of tetR, followed by *V4Tc, V2Tc* and *V3Tc* (Fig. 5B). Incorporation of the tetR repressor gene under the control of the PJ23119 promoter silenced all the synthetic *tet* promoters, as in *E. coli* (Fig. 5B). Addition of aTc activated the four *tet* promoters and remarkably, the induced *V2Tc* promoter showed the highest mCardinal induction exceeding its constitutive expression by ∼10-fold and matching the *tacI* promoter. The *V2Tc* promoter was revealed to be ∼2 and 3-fold more efficient than the *V1Tc* and *V4Tc* promoters after induction respectively, while the *V3Tc* proved to be inefficient in *P. putida* (Fig. 5C). These results highlight the importance of the consensus sequences targeted by σ^70^ to amplify the strength of inducible promoters also in *P. putida*.

### Strength and regulation of synthetic *lac* and *tet* promoters in *Vibrio natriegens*

To further validate the hypothesis that the incorporation of the σ^70^ consensus sequences improve the strength of inducible promoters in various gram-negative bacteria, we evaluated our circuits in the marine bacterium *V. natriegens* which has gained popularity for routine molecular biology applications due its ability to double in <10 min [20, 21] (Fig. 6). In this host, previous studies indicated that the *tacI* promoter produced GFP upon activation with IPTG, while induction of the *tet* promoter resulted in low GFP yields [21]. The constitutive and inducible versions of the synthetic *lac* and *tet* promoters were evaluated in the pColE1 derived plasmid with ∼300 copies per cell [21], except for the constitutive *V2Lac*, which was evaluated in pCloDF13 derived vector with ∼64 copies per cell in this strain [42]. The constitutive *V2Lac* (Fig. 6A) and *V4Tc* (Fig. 6B) promoters outperformed *tacI* by ∼16 and ∼2-fold respectively (Fig. 6C), while *V3Lac, V1Tc* and *V2Tc* produced similar levels as *tacI. V1Lac* and *V3Lac* underperformed *tacI* by ∼2.7-fold, and *V3Tc* showed no activity (Fig. 6C). Incorporation of the transcriptional regulators lacI and tetR turned OFF all the synthetic promoters except for *V2Lac*, which continued to show leakage, as in *E. coli* and *P. putida* (Fig. 6A). Induction of the synthetic *lac* promoters by IPTG revealed that only the *V2Lac* promoter was fully activated outpassing its constitutive version by ∼1.3-fold and showed a potential dynamic range of ∼25-fold (Fig. 6A,C). The *V3Lac* reached 30% of its full potential, and the *V1Lac* and *V4Lac* showed no induction (Fig. 6A). While addition of aTc activated all four synthetic *tet* promoters, the *V3Tc* and *V4Tc* promoters rather showed weak mCardinal production, and *V1Tc* produced 2-fold more mCardinal than its constitutive variant (Fig. 6B). As predicted, the *V2Tc* promoter, which harbors the 2 consensus σ^70^ boxes, demonstrated strong inducibility producing 11-fold more mCardinal than the *tacI* promoter and displaying full potential of its dynamic range producing 100-fold more mCardinal than its OFF version (Fig. 6B,C). These results confirmed that the adaptation of the two consensus boxes recognized by the σ^70^ in the *V2lac* and *V2Tc* promoters improved the performance of the inducible *lac* and *tet* promoters in also *V. natriegens*.

**Figure 6.**
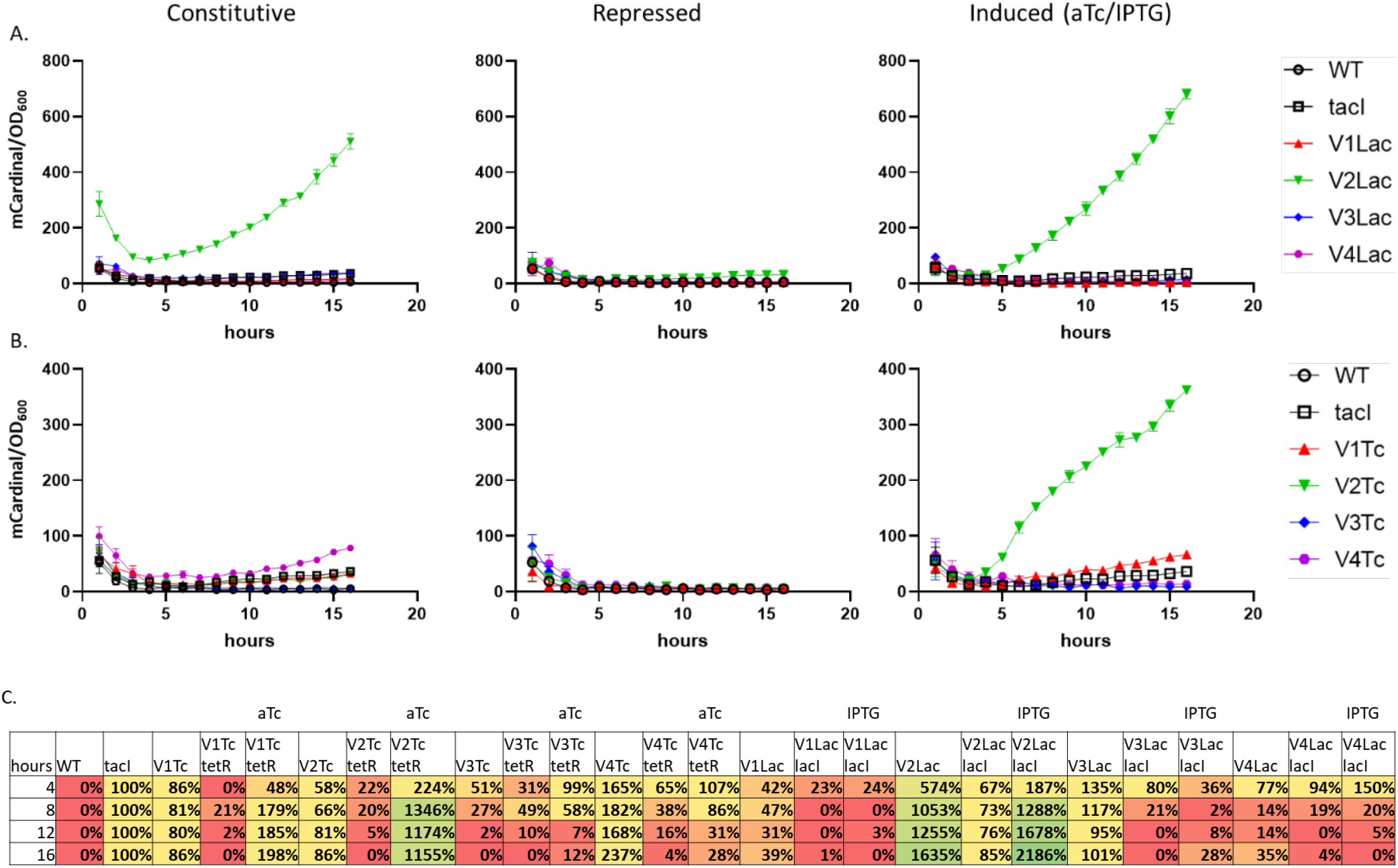
Time courses of mCardinal production in *V. natriegens*. Synthetic *lac* **(A)** and *tet* (**B**) promoters in their constitutive, repressed and induced states. **(C)** Direct comparison of each promoter under evaluation against the constitutive promoter *tacI* in *V. natriegens. V. natriegens* carrying *tacI-mCardinal* was normalized to 100% mCardinal production and the wild type *V. natriegens* strain normalized to 0% mCardinal production. For all samples, the fluorescence mean was normalized by the cell density (OD_600_). N=4. Error bars +/-SD.

## Discussion

Production of recombinant proteins is and will continue to be one of the main tools scientists use to understand biological processes and transfer academic results to industrial applications. Over the last 30 years, the pET system alone allowed the production of >220,000 recombinant proteins [1]. Despite the astonishing number of proteins produced by the pET system, its failure rate is impossible to determine. The pET system is well known to induce the formation of inclusion bodies, a major drawback in the production of soluble proteins. pET requires specific strains that carry the T7 RNA polymerase, it lacks real tuneability, and fails to keep the target gene OFF. Leakiness in the OFF state negates the main advantage of an inducible system, which is intended to permit time-or context-specific control of gene expression. Our results show that inclusion of the σ70 consensus sequences into the *lac* and *tet* inducible promoters improve both repression and induction. The tight transcriptional control does not require any particular strain background, and permits rapid expression of soluble proteins, including toxic proteins such as CocE. We anticipate that our solution offers a significant advance to the biotechnology industry in offering additional platforms for exogenous protein expression and purification.

Not even a perfect expression system will allow *E. coli* to produce all proteins. Therefore, complementary expression hosts could provide alternative solutions. We have demonstrated our inducible promoters can be easily optimized to be used in a variety of gram-negative hosts, including *P. putida* and *V. natriegens*, thus widening the applicability of these tools to a broad spectrum of bacteria.

As with the original *lac, tet*, and pET systems, the synthetic promoters presented here cannot be tuned by the inducer concentration. However, different yields of recombinant protein can be achieved based on promoter and host selection. Overall, the incorporation of the consensus −35 sequence and Pribnow (−10) box unlocks the strength of the *lac* and *tet* promoters in gram-negative bacteria, facilitating the production of any given target gene in different host with the same set of plasmids.

## Methods

### Bacterial strains

*E. coli* DH10B, *P. putida* AG4775 derivative of JE90 [41], and V. natriegens Vmax X2 (Codex DNA, Inc.) were used to evaluate the synthetic *lac* and *tet* promoters. The plasmid pET21a-mCardinal was evaluated in BL21 strain. Selective markers kanamycin, ampicillin, spectinomycin and streptomycin were supplemented to LB medium when required. *E. coli* strains were transformed by electroporation and *V. natriegens* by chemical transformation. Integration of plasmids into *P. putida* chromosome was performed by electroporation following the protocol as described [41]. Plasmid information is listed in Supplementary Table S1.

### Molecular cloning

Polymerase chain reactions (PCR) were carried out with Phusion DNA polymerase (ThermoFisher Scientific, USA). Digestion of DNA was performed with fast digest restriction enzymes and DNA fragments were join by T4 DNA ligase or Gibson assembly kit (ThermoFisher Scientific, USA). Oligonucleotides and DNA synthesis were ordered to IDT (IDT, USA). DNA sequencing was performed by QuintaraBio (QuintaraBio, USA). Shuttle vectors pJH0204 and pJH0228 were kindly provided by Dr. Adam Guss, Oak Ridge National Laboratory [41]. The synthetic *Vlac* and *VTc* promoters were synthesized as gBLocks and incorporated into the MCS of pJH0204 using BamHI and XhoI. Further, sfGFP and mCardinal ORF were codon optimized for *P. putida* and inserted downstream the synthetic promoters using NdeI and EcoRI. The transcriptional regulator tetR was synthesized together with the pJ23119 promoter and codon optimized for *P. putida* and inserted after mCardinal with EcoRI and XhoI. LacI was PCR amplified from pET21a vector and placed under the control of pJ23119 using NcoI and XhoI, but this combination resulted toxic for *E. coli*, therefore lacI was PCR amplified together with its native promoter and ubicated after mCardinal using EcoRI and XhoI. pJH0228-V2lac-mCardinal and pJH0228-V2lac/lacI-mCardinal vectors were produced using the same cloning strategy. mCardinal was inserted into pET21a using NdeI and XhoI. The transcriptional unit V2Tc/tetR-mCardinal was inserted into the pUC19 vector by Gibson removing the NdeI RE nucleotide sequence in the plasmid and incorporating terminators insulating the transcriptional unit. DNA sequence of cocE was codon optimized for *P. putida* and cloned into the pET21a and *V2Tc-tetR* expression system using NdeI and HindIII. Primers used in this study are listed in Supplementary Table S2. Sequences of cocE, tetR, and mCardinal, codon optimized for *P. putida*, and the pJ23119 promoter sequence, are available in the Supplementary Information.

### Bacterial growth curve and sfGFP/mCardinal measurements with plate reader

Growth curves and fluorescence measurements were performed in 96-well clear bottom microplates (Corning Incorporated, NY) using the BioTek Synergy H1 Hybrid Multi-Mode Reader. Briefly, ON cultures of the recombinant strains were diluted into fresh LB medium to reach OD_600_ 0.05. *E. coli* experiments were performed at 37°C, and *P. putida* and *V. natriegens* at 30°C, always at 355 cpm. OD_600_ and fluorescence (sfGFP excitation = 485 & emission = 510; mCardinal excitation = 604 & emission = 659) were monitored every hour for 12 hours. *E. coli* cultures were induced at 3 hours with 0.2 mM IPTG and/or 0.1 µg/mL aTc, *P. putida* at 3 hours with 5 mM IPTG and/or 1 µg/mL aTc, and *V. natriegens* at 2 hours with 0.2 mM IPTG and/or 0.1 µg/mL aTc when required.

### SDS-PAGE

SDS-PAGE was carried out on a 4–12% Bis-tris Midi Protein Gel in an XCell4 SureLock Midi system (Invitrogen, USA). *E. coli, P. putida* and *V. natriegens* cell extracts were obtained from 10 mL LB culture inoculated with 2.5% ON culture and induced at ∼OD_600_ 700 with 0.2 mM IPTG or 0.1 µg/mL aTc accordingly. Induced cultures were grown for 5 hours, and 2 mL culture were spun down at 14.000 rpm and 4°C for 10 minutes and frozen at -20°C for further analysis. Total protein concentration was estimated with Thermo Scientific NanoDrop one and ∼10 mg total protein of each cell extract was loaded into the SDS-gel.

### Expression of CocE and measurement of benzoic acid production

Recombinant *E. coli* BL21 strain carrying the pET21a-cocE and DH10B containing either p*σ*^*70*^ V2TcR-cocE or p*σ*^*70*^ V2TcR-cocE19 plasmids were cultivated overnight at 30° C at 220 rpm and fresh cultures were started with 5% of the ON culture and induced at ∼OD_600_ 700 with 0.2 mM IPTG or 0.1 µg/mL aTc accordingly. 1 mL samples were collected each hour for 6 hours by centrifugation at 4° C and 14000 rpm and stored at -80° C. Cell pellets were disrupted using 150 µL of B-PE Complete Bacterial Protein Extraction Reagent (ThermoFisher Scientific) for 25 minutes at room temperature and soluble fractions were collected after centrifugation for 25 minutes at 4° C and 14000 rpm. ∼ 3 mg/mL of soluble fractions were incubated with 0.015 mg of cocaine for 20 minutes at 28° C and benzoic acid production was estimated using the benzoic acid detection kit for feed (Attogene, EZ2013-03).

## Competing interests

The authors have filed a provisional patent application for these promoter designs.

## Supplementary Information

A document containing Supplementary Figures S1-S3, Supplementary Tables S1 and S2, and additional sequence information accompanies this manuscript.

## Data sharing plan

Raw datasets for all experiments are available by request to N.G.F. All reagents are available upon request to N.G.F.

## Acknowledgements

We thank Dr. Adam Guss of Oak Ridge National Laboratory for the gift of pJ0204, pJ0228, and *P. putida* strains used in this work. We thank Dr. Eric Young for sharing equipment essential to the success of this work. This work was supported by new faculty start-up funds from Worcester Polytechnic Institute to N.G.F.

## Notes

### Competing Interest Statement

The authors have filed a provisional patent application based on the promoter designs reported in this manuscript.

