## Supplementary information for "Unlocking the strength of inducible promoters in gram-negative bacteria"

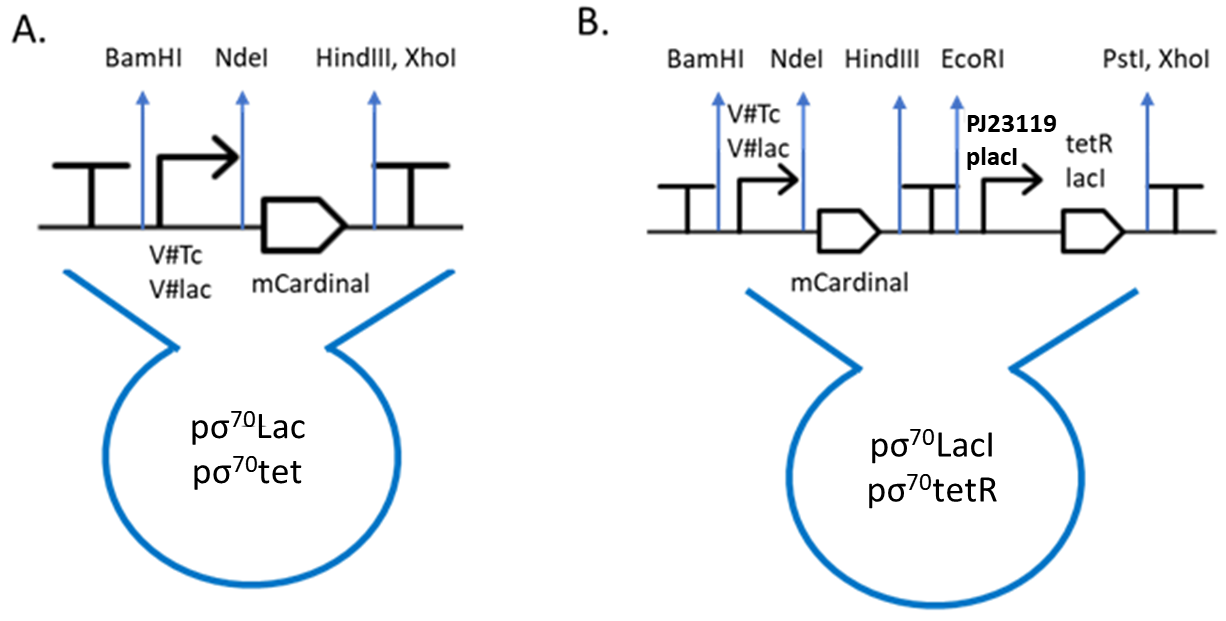

**Figure S1.** Genetic organization of the plasmids produced in this study. **A.** Constitutive version of the synthetic *lac* and *tet* expression systems, and **B.** Inducible versions of the synthetic *lac* and *tet* expression systems in pColE1 backbone. Plasmids contain the Kanamycin selection marker and the attP sites specific fox Bxb1 integrase.

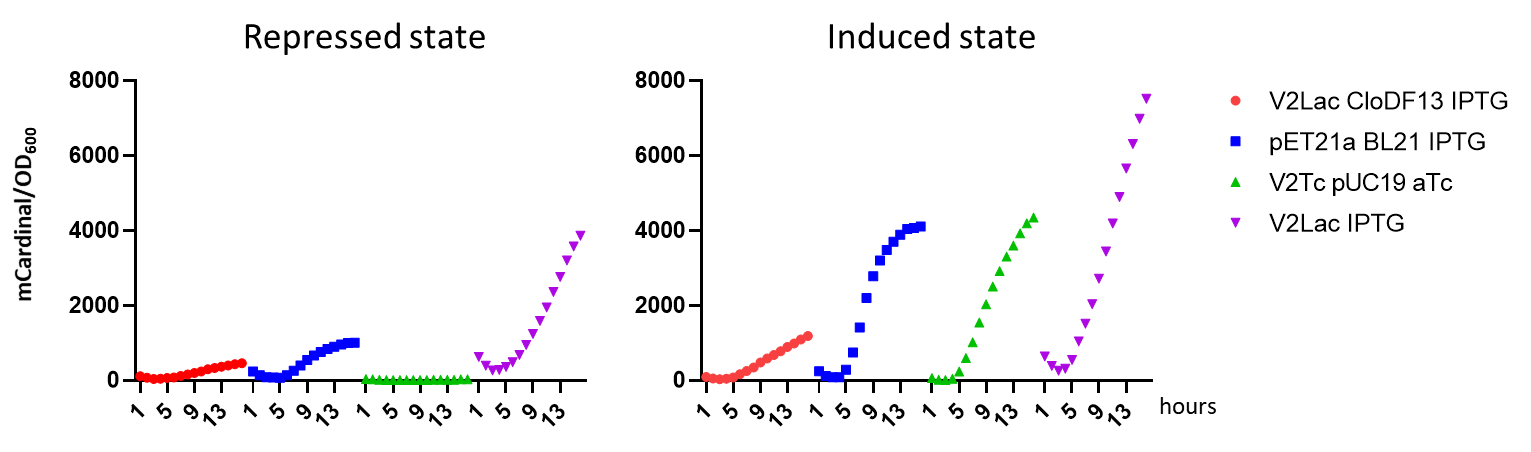

**Figure S2.** Comparison of the pET21a expression system in *E. coli* BL21 versus the synthetic *V2Lac/lacI* and *V2Tc/tetR* promoters in *E. coli* DH10B. Cultures were induced after 3 hours of cultivation and mCardinal mean was normalized by the cell density (OD_600_). N=4. Error bars +/- SD.

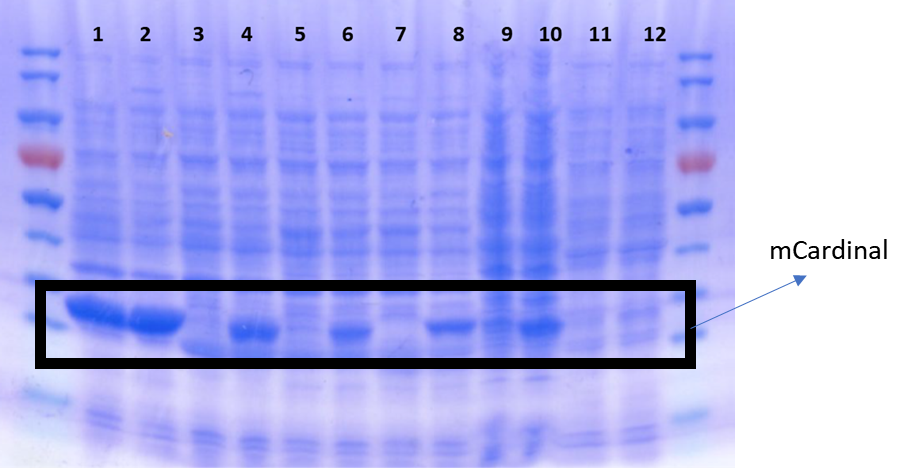

**Figure S3.** SDS gel of recombinant *E. coli*, *P. putida* and *V. natriegens* strains induced at OD_600_ 0.7 with IPTG/aTc and cultured for 5 hours.

1. *E. coli* BL21 pET21a-mCardinal
2. *E. coli* BL21 pET21a-mCardinal 0.2 mM IPTG
3. *E. coli* BL21 pJH0204 V3Lac-mCardinal/placI-lacI
4. *E. coli* BL21 pσ^70^ V3LacI-cardi 0.2 mM IPTG
5. *E. coli* DH10B pσ^70^ V3LacI-cardi
6. *E. coli* DH10B pσ^70^ V3LacI-cardi 0.2 mM IPTG
7. E. coli DH10B pσ^70^ V2TcR-cardi
8. *E. coli* DH10B pσ^70^ V2TcR-cardi 0.1 ug/mL Tc
9. *V. natriegens* pσ^70^ V2TcR-cardi 0.1 ug/mL Tc
10. *V. natriegens* pσ^70^ V2LacI-cardi 0.2 mM IPTG
11. *P. puitda* pσ^70^ V2TcR-cardi 0.1 ug/mL Tc
12. *P. puitda* pσ^70^ V2LacI-cardi 0.2 mM IPTG

**Table S1.** List of vectors generated in this study

| **Name** | **backbone** | **Origin of replication** | **Selective marker** | **Construct** | **Origin** |
| --- | --- | --- | --- | --- | --- |
| pJH0204 | pJH0204 | pColE1 | Kanamycin |  | Gift Dr. Adam Guss, ORNL |
| pJH0228 | pJH0228 | pCloDF13 | Spectinomycin/Streptomycin |  | Gift Dr. Adam Guss, ORNL |
| pJH0204F | pJH0204 | pColE1 | Kanamycin |  | This study |
| pFCtacI-GFP | pJH0204 | pColE1 | Kanamycin | *tacI*-sfGFP | This study |
| pFCtacI-cardi | pJH0204 | pColE1 | Kanamycin | *tacI*-mCardinal | This study |
| pσ^70^ V1Lac-cardi | pJH0204 | pColE1 | Kanamycin | *V1lac*-mCardinal | This study |
| pσ^70^ V2Lac-cardi13 | pJH0228 | pCloDF13 | Spectinomycin/Streptomycin | *V2lac*-mCardinal | This study |
| pσ^70^ V3Lac-cardi | pJH0204 | pColE1 | Kanamycin | *V3lac*-mCardinal | This study |
| pσ^70^ V4Lac-cardi | pJH0204 | pColE1 | Kanamycin | *V4lac*-mCardinal | This study |
| pσ^70^ V1LacI-cardi | pJH0204 | pColE1 | Kanamycin | *V1lac/lacI*-mCardinal | This study |
| pσ^70^ V2LacI-cardi | pJH0204 | pColE1 | Kanamycin | *V2lac/lacI*-mCardinal | This study |
| pσ^70^ V2Lac-cardi13 | pJH0204 | pCloDF13 | Spectinomycin/Streptomycin | *V2lac/lacI*-mCardinal | This study |
| pσ^70^ V3LacI-cardi | pJH0204 | pColE1 | Kanamycin | *V3lac/lacI*-mCardinal | This study |
| pσ^70^ V4LacI-cardi | pJH0204 | pColE1 | Kanamycin | *V4lac/lacI*-mCardinal | This study |
| pσ^70^ V1Tc-cardi | pJH0204 | pColE1 | Kanamycin | *V1Tc*-mCardinal | This study |
| pσ^70^ V2Tc-cardi | pJH0204 | pColE1 | Kanamycin | *V2Tc*-mCardinal | This study |
| pσ^70^ V3Tc-cardi | pJH0204 | pColE1 | Kanamycin | *V3Tc*-mCardinal | This study |
| pσ^70^ V4Tc-cardi | pJH0204 | pColE1 | Kanamycin | *V4Tc*-mCardinal | This study |
| pσ^70^ V1TcR-cardi | pJH0204 | pColE1 | Kanamycin | *V1Tc/tetR*-mCardinal | This study |
| pσ^70^ V2TcR-cardi | pJH0204 | pColE1 | Kanamycin | *V2Tc/tetR*-mCardinal | This study |
| pσ^70^ V3TcR-cardi | pJH0204 | pColE1 | Kanamycin | *V3Tc/tetR*-mCardinal | This study |
| pσ^70^ V4TcR-cardi | pJH0204 | pColE1 | Kanamycin | *V4Tc/tetR*-mCardinal | This study |
| pσ^70^ V2TcR-cardi19 | pUC19 | pMB1 | Ampicillin | *V2Tc/tetR*-mCardinal | This study |
| pσ^70^ V2TcR-cocE | pJH0204 | pColE1 | Kanamycin | *V2Tc/tetR*-cocE | This study |
| pσ^70^ V2TcR-cocE19 | pUC19 | pMB1 | Ampicillin | *V2Tc/tetR*-cocE | This study |
| pET21a-cardi | pET21a | pET | Ampicillin | *Pet*-mCardinal | This study |
| pET21a-cocE | pET21a | pET | Ampicillin | *pET-*cocE | This study |

**Table S2.** List of primers generated in this study

| **Primer name** | **Sequence 5’ to 3’** | **logic** |
| --- | --- | --- |
| pUC19 modified FW XhoI | attgcCTCGAGACTGATTTTTAAGGCGACTGATGAGTCGCCTTTTTTTTGTCTAGCTAACTCACATTAATTGCGTTGCGCT | Incorporate terminators into pUC19 vector and BamHI/XhoI sites |
| pUC19 modified FW XhoI | gcaatGGATCCagtcaaaagcctccgaccggaggcttttgactAGTACAATCTGCTCTGATGCCGCATAGTT |  |
| 204F1_fwd | Ggcgcgtactccaaaaggatctaggtgaagatc | Incorporate BamHI, NdeI, HindIII, KpnI, EcorI, PstI, XbaI, NcoI, XhoI RE into the MCS of pJH0204 and remove NcoI from kanamycin cassette |
| 204F1_rev | gagttcttctgattagacaaaaaaaaggcgac |  |
| 204F2_fwd | tttttttgtctaatcagaagaactcgtcaagaagg |  |
| 204F2_rev | gcccgacggcgaggatctcgtcgtgacgcatg |  |
| 204F3_fwd | cacgacgagatcctcgccgtcgggcatccgc |  |
| 204F3_rev | gtctagactgcaggaattcaagcttcatatgggatccggaccaaaacgAAA |  |
| 204F4_fwd | aagcttgaattcctgcagtctagaccatggctcgaggacgaacaataaggc |  |
| 204F4_rev | Cctagatccttttggagtacgcgcccgggga |  |
| placI upstream | TACTGGTTTCACATTCACCAC | Sequencing primers |
| lacI downstream | GTGGTGAATGTGAAACCAGTAA |  |
| lacI upstream | TTTCCAGTCGGGAAACCT |  |
| lacI downstream2 | AGGTTTCCCGACTGGAAA |  |
| tetR upstream | CCAATACAACGTGGGTTGCT |  |

DNA sequence of mCardinal codon optimized for *P. putida*

ATGGTGAGTAAGGGTGAGGAGCTCATTAAGGAGAACATGCACATGAAGCTGTATATGGAGGGCACCGTAAACAACCACCACTTCAAGTGTACCACCGAGGGTGAAGGTAAACCCTACGAGGGGACGCAGACCCAACGCATCAAGGTCGTGGAGGGCGGCCCGCTGCCTTTCGCATTCGACATTCTGGCGACCTGTTTTATGTACGGCTCGAAGACCTTCATCAACCACACCCAAGGCATCCCGGACTTCTTCAAGCAGAGCTTCCCTGAGGGCTTCACCTGGGAGCGCGTCACCACGTATGAAGACGGTGGGGTGCTCACCGTGACCCAGGACACGAGCTTGCAGGATGGCTGCTTGATTTACAACGTCAAGCTGCGCGGGGTGAACTTCCCTAGCAACGGGCCAGTGATGCAGAAAAAGACGCTGGGTTGGGAGGCCACCACCGAGACCCTGTACCCGGCCGACGGGGGGCTGGAAGGGCGGTGCGATATGGCCCTGAAATTGGTCGGCGGCGGTCATTTGCACTGCAATCTCAAGACCACGTACCGCTCCAAGAAACCCGCCAAAAACCTGAAGATGCCTGGTGTTTATTTTGTCGACCGGCGCCTGGAGCGCATCAAGGAAGCGGACAATGAGACGTACGTGGAACAGCACGAAGTGGCCGTGGCTCGTTATTGCGATCTGCCGTCGAAGCTGGGTCACAAACTGAACGGCATGGATGAGCTGTACAAAGATTATAAGGATGATGACGACAAGTAA

DNA sequence of tetR codon optimized for *P. putida*

ATGTCCCGCCTGGATAAATCGAAAGTGATTAACTCGGCCCTCGAATTGCTGAATGAAGTCGGTATCGAGGGGCTGACGACCCGTAAATTGGCACAAAAGTTGGGGGTGGAGCAACCCACGTTGTATTGGCACGTCAAAAATAAGCGGGCATTGCTGGATGCCCTCGCTATTGAAATGTTGGATCGCCACCATACCCATTTCTGTCCACTGGAGGGCGAGTCCTGGCAGGACTTTCTCCGCAACAACGCGAAATCCTTTCGCTGTGCACTCTTGTCCCATCGGGACGGTGCTAAGGTGCACTTGGGCACCCGTCCCACCGAAAAACAATACGAAACCTTGGAAAATCAATTGGCGTTTTTGTGCCAGCAAGGGTTTAGCTTGGAGAATGCTCTCTATGCGCTCTCGGCTGTCGGGCACTTTACGTTGGGGTGCGTGTTGGAGGACCAGGAGCATCAAGTCGCAAAAGAGGAGCGTGAAACCCCAACCACGGACTCGATGCCACCTCTGCTCCGCCAAGCTATCGAACTCTTCGATCATCAGGGCGCGGAGCCAGCCTTCCTCTTTGGGCTGGAGCTGATTATCTGCGGTTTGGAAAAACAACTCAAGTGTGAAAGCGGGTCCTAA

DNA sequence of cocE codon optimized for *P. putida*

CATATGGTGGACGGTAATTATTCGGTAGCGTCCAACGTTATGGTGCCGATGCGCGACGGGGTGCGCTTGGCTGTAGATCTGTACCGCCCGGACGCAGATGGCCCTGTACCGGTCCTGCTGGTCCGCAACCCCTACGACAAATTCGACGTGTTCGCTTGGAGTACGCAGAGCACGAACTGGCTGGAATTTGTGCGCGATGGGTACGCCGTCGTCATCCAAGACACCCGGGGCCTCTTTGCATCCGAAGGTGAGTTCGTTCCACATGTTGATGACGAGGCGGATGCGGAAGACACGCTGAGCTGGATCTTGGAACAAGCATGGTGCGACGGCAATGTGGGTATGTTCGGTGTAAGCTACCTGGGCGTTACGCAGTGGCAAGCTGCTGTTAGCGGTGTGGGTGGTTTGAAGGCAATCGCCCCGAGCATGGCGAGCGCGGATCTGTACCGTGCCCCCTGGTACGGTCCTGGCGGCGCCCTGAGCGTGGAAGCACTCCTGGGCTGGAGCGCATTGATCGGTACGGGCCTGATTACCAGCCGTAGCGATGCCCGCCCGGAAGACGCAGCCGACTTCGTACAGCTGGCAGCCATCCTGAACGATGTGGCCGGTGCCGCAAGCGTGACCCCTCTGGCCGAACAGCCCTTGTTGGGCCGCCTGATCCCTTGGGTGATCGACCAGGTGGTGGACCATCCAGACAACGACGAGTCGTGGCAGAGCATCTCGCTCTTTGAACGTTTGGGTGGGCTCGCTACCCCGGCCTTGATTACCGCCGGGTGGTACGATGGCTTCGTGGGCGAGAGCCTCCGTACCTTCGTAGCTGTGAAGGACAACGCGGATGCGCGTCTGGTGGTGGGGCCGTGGAGCCACAGCAATCTGACCGGCCGTAATGCCGACCGTAAGTTTGGGATCGCCGCGACCTACCCCATCCAGGAGGCGACGACCATGCACAAGGCTTTTTTCGACCGGCACCTCCGTGGCGAGACCGATGCCCTGGCAGGGGTGCCCAAGGTGCGCCTCTTCGTAATGGGTATCGATGAGTGGCGCGACGAGACCGACTGGCCATTGCCAGATACCGCTTACACGCCTTTTTACCTCGGGGGCTCCGGTGCGGCCAACACGAGCACGGGTGGTGGGACCCTGTCGACCTCGATCAGCGGCACGGAGTCGGCGGACACCTACCTGTATGATCCTGCCGACCCCGTGCCAAGTCTGGGCGGCACCCTCCTCTTCCATAATGGGGACAACGGTCCAGCTGACCAGCGCCCGATTCACGATCGCGACGACGTGCTGTGCTACTCCACCGAGGTGTTGACCGACCCCGTGGAAGTAACGGGGACGGTTTCGGCTCGCCTGTTCGTGTCCTCGTCGGCCGTGGATACCGATTTTACCGCCAAGTTGGTCGACGTGTTCCCCGATGGTCGGGCAATCGCTCTCTGCGACGGCATCGTGCGTATGCGCTACCGGGAGACCTTGGTAAATCCTACGCTCATTGAGGCCGGTGAGATTTACGAGGTGGCTATTGATATGCTGGCCACCAGCAACGTGTTTTTGCCGGGCCACCGCATCATGGTGCAAGTTAGCAGCTCGAACTTCCCGAAGTACGACCGCAACTCCAACACCGGCGGCGTCATCGCTCGCGAGCAACTGGAGGAAATGTGCACCGCCGTAAACCGCATTCACCGCGGCCCCGAACACCCGTCCCATATCGTGCTGCCGATCATTAAGCGCGACTATAAGGACGACGACGATAAGTGAAAGCTT

DNA sequence of pJ23119 promoter

TTGACAGCTAGCTCAGTCCTAGGTATAATGCTAGCCGCAGTAAGAGAGGAATGTACAC
